# Hierarchical Chromatin Rewiring Orchestrates Early Transcriptional Regulation in Mouse Cerebral Cortex Following Focal Ischemia

**DOI:** 10.64898/2026.01.28.702387

**Authors:** Hadjer Namous, Raghu Vemuganti

## Abstract

Ischemic stroke triggers massive transcriptional reprogramming, yet how the brain’s higher-order chromatin architecture orchestrates this response remains unknown. We mapped the spatiotemporal reorganization of the genome in the mouse peri-infarct cerebral cortex following transient middle cerebral artery occlusion at 6h and 24h of reperfusion. By integrating high-resolution Hi-C data with transcriptomic and cis-regulatory landscapes, we show that stroke induces a hierarchical rewiring of genome architecture across compartments, domains, and loops. Early A to B compartment shifts were largely transcriptionally silent for coding genes, whereas B compartments were enriched for upregulated noncoding RNAs. We also observe structural dependencies between scales. Gained loops do not independently drive differential expression. Instead, their regulatory potential is gated by their domain context. Loops nested within expanded Topologically Associating Domains (TADs) show a higher percentage of stroke-responsive transcripts. Flow analyses indicate that gained TADs establish the primary scaffold for transcriptional responses, while compartment identity refines the specificity of noncoding RNA regulation. These findings suggest that post-stroke gene expression follows a selective, multi-scale architectural hierarchy, with chromatin remodeling as a central regulator of the early ischemic stress response and genome architecture is a determinant of transcriptional outcomes.

## Introduction

The mammalian genome is folded in a hierarchical three-dimensional (3D) architecture that spans multiple scales to regulate gene expression, cellular identity and dynamic responses to environmental cues.^1–3^ Fine-resolution chromatin loops connect regulatory elements such as promoters, enhancers and super-enhancers (SEs).^2–5^ In addition, topologically associating domains (TADs) form insulated neighborhoods that constrain regulatory element communication.^6–8^ At a broader scale, A (active) and B (inactive) compartments reflect transcriptional activity and chromatin state.^3,9,10^ Together, these layers form the spatial organization of the genome, that dynamically coordinates transcription across time and cellular states.

Reorganization of genome topology is a hallmark of stress response across biological systems. Chromatin loops can form de novo following immune activation or oncogenic signaling,^23,24^ and TAD boundaries and compartments alter in response to DNA damage and cellular differentiation.^25,26^ In the brain, chromatin remodeling has been observed during development and neurodegeneration.^11–13^ Recent studies showed altered cerebral loops and TAD reorganization following experimental stroke in rodents.^27^ However, the temporal dynamics and multi-scale coordination of these changes in the post-stroke brain have not been explored yet.

We presently conducted a multi-scale, time-resolved mapping of the spatiotemporal genome architecture in the adult mouse cerebral cortex following transient middle cerebral artery occlusion (MCAO). Chromatin remodeling across compartments, TADs, and loops was evaluated by integrating high-resolution Hi-C data with transcriptomic and cis-regulatory annotations, including promoters, enhancers, CTCF sites, and SEs at 6h and 24h reperfusion following transient MCAO. The structural transitions were linked to post-ischemic transcriptional outcomes.

## Methods

### Transient focal ischemia

All procedures were approved by the University of Wisconsin Research Animal Resources and Care Committee and conformed to the *Guide for the Care and Use of Laboratory Animals (8th ed., 2011)*. Adult male C57BL/6J mice (27 ± 2 g; Jackson Laboratory) were subjected to transient MCAO using a silicon-coated 6-0 monofilament (Doccol) under isoflurane anesthesia as previously described.^28^ Core temperature was kept at 37°C with a feedback-controlled heating pad. Following 1h of occlusion, cohorts of mice were euthanized at 6h and 24h of reperfusion. Sham-operated mice underwent the same procedures without occlusion. Peri-infarct cortical tissue and corresponding sham tissue were dissected, flash-frozen in liquid nitrogen. 36 mice were used to generate n = 4 pooled biological replicates/condition (sham, 6h reperfusion and 24h reperfusion), with each replicate comprising 3 randomly assigned animals.

### Hi-C library preparation

Hi-C libraries were generated using the Arima-HiC kit (A510008) and the Arima Library Prep Module (A303011) following manufacturer’s protocols. Briefly, tissue was crosslinked, pulverized using liquid nitrogen, digested with the Arima enzyme cocktail, proximity-ligated, reverse-crosslinked and purified. DNA (∼3 µg) was fragmented to 200–600 bp, and 200 ng of biotin-enriched DNA was size-selected, indexed and amplified. Libraries were sequenced on an Illumina NovaSeqPlus platform (150 bp paired-end), with a target of ≥ 600 million reads per library.

### Hi-C processing and feature calling

Raw FASTQ files were quality checked using MultiQC and processed using Juicer v1.6 (BWA-MEM mapping to mm39/GRCm39, MAPQ ≥ 30, duplicates removed and KR normalization applied). Three sample libraries were retained for each group based on the library QC quality (see Supplementary Table S1). Contact matrices were generated at multiple resolutions. Genome features were called using Juicer Tools v2.20.00 on KR-normalized matrices as described below. To identify compartments, eigenvectors were calculated via principal component analysis (PCA) at 1Mb resolution. Bins overlapping centromeres or telomeres were removed. Z-score normalization was performed per-sample eigenvectors. To harmonize compartment calls across samples, A/B labels were oriented by PC1 sign, designating bins positively correlated with gene density as compartment A. Compartment bins were considered Stable (A or B) if they retained their sham label. A to B and B to A bins were defined as switching relative to sham. TADs were called per replicate using Juicer/Arrowhead at 10 kb resolution. Domains were defined from left-boundary start to right-boundary end. Interchromosomal pairs were excluded, and intervals overlapping annotated centromeres/telomeres were removed with harmonized chromosome naming. Within each group (sham, 6h reperfusion, and 24h reperfusion), reproducible TADs were defined as those present in all replicates within a ±20 kb boundary tolerance. TADs were compared to sham using reciprocal overlap (RO ≥ 0.8). Domains with boundary displacement of ≤10 kb at both ends were labeled stable; those with >10 kb shift at either end were labeled shifted. Domains found only at 6h reperfusion or 24h reperfusion were classified as gained, and those found only in sham were called lost. For fusion/fission detection, a broader ±40 kb tolerance was used: a single condition TAD overlapping multiple sham TADs was considered merged, and a single sham TAD overlapping multiple condition TADs was considered split. Category2 labels (merged/split) were only assigned to TADs already labeled as gained or lost to avoid misclassifying boundary jitter as architectural reorganization. The 10 kb resolution is consistent with high-depth Hi-C standards, and the ±10/±40 kb thresholds reflect the bin size and expected uncertainty in boundary calls. TAD boundary strength was quantified using the KR-normalized z-scores provided by Arrowhead. Boundaries with z-scores above 1 were considered strong, while lower scores indicate weaker insulation. Chromatin loops were detected at 10 kb resolution using HiCCUPS. Within each condition, loops were defined as reproducible if they passed HiCCUPS significance in all biological replicates, where a loop “passed” if adj.p was <0.05 and the corresponding bin count was ≥3 in any context (BL, Donut, H, or V). Reproducible loops were further classified by strength and length by defining them as strong (observed contacts ≥15, obs/exp ≥ 2.5 in any context, adj.p <0.05) or weak (observed contacts <10 and obs/exp<1.5 across all four contexts). Loops are considered as short (<200 kb) and long (≥200 kb). Gained and lost loops were defined relative to sham by comparing the union of reproducible intra-chromosomal loops across timepoints. Each unique loop (based on exact anchor coordinates) was checked for presence or absence in sham, 6h reperfusion and 24h reperfusion. A loop was classified as gained if present at a reperfusion time point, but absent in sham, and lost if present in sham, but absent at both reperfusion time points.

### Transcriptome profiling and differential expression

We used the transcriptome data previously generated by our group that profiled both noncoding RNAs and coding RNAs using Arraystar microarray platform.^28^ Differential expression was assessed at the transcript level using GRCm39 by converting the original transcript IDs and their respective genomic coordinate from GRCm38.

### Regulatory annotation of cis elements

Candidate cis-regulatory elements (cCREs), including enhancers, promoters, CTCF-binding sites, and SEs were obtained from the WengLab SCREEN database for mouse (https://screen.wenglab.org/about] (https://screen.wenglab.org/about; mm10/GRCm38). Stroke-specific SEs were derived from SEdb mouse brain atlas (http://www.licpathway.net:8081/sedb/download\mm.php). All blacklisted regions such as centromeres and telomeres were excluded. As the regulatory annotations were originally in mm10/GRCm38 coordinates, we used Ensembl Assembly Converter using mouse as species to lift over all coordinates to mm39/GRCm39 (https://useast.ensembl.org/Homo\sapiens/Tools/AssemblyConverter).

### Integration of 3D features with gene expression

Loop anchors (anchor 1 and anchor 2) were overlapped with cis-regulatory annotations (promoters, enhancers, CTCF sites, and SEs), and per-loop element counts and densities were calculated. Associated transcripts were assigned to loops by intersecting loci of both anchors with regulatory elements and transcript annotations. Transcript-level differential expression data were then linked to loop, TAD, and compartment contexts. Loops were classified as inside-TAD, cross-boundary, or outside by intersecting anchor coordinates with TADs.

### Statistical analysis

Unless otherwise noted, all statistical tests were two-sided and p values were adjusted using Benjamini-Hochberg false discovery rate for adjusted p value (adj.p). For enrichment analysis, Fisher’s exact test was used and odds ratio (OR) with >95% confidence interval was reported. Distributions (e.g., boundary strength, loop length and regulatory element density) were compared using the Wilcoxon rank-sum test. Statistical analyses were performed in R (version 4.4.1). Detailed package and session information is provided in the Supplementary Information.

### Data and code availability

Code for raw and processed Hi-C and transcriptome data analysis is available upon reasonable request.

## Results

### Genome-wide compartment switching after stroke is rapid and reversible

A/B compartment analysis at 6h and 24h reperfusion after focal ischemia showed pronounced chromatin reorganization compared with sham (Fig. 1A). The proportion of bins undergoing compartment shift was significantly higher at 6h than 24h (58% vs 51%, p<0.01), indicating greater compartmental flux immediately after ischemia. This was driven by A to B transitions (29% at 6h; 25% at 24h; p<0.01), while B to A changes were not significant. Stable A compartments became significantly more frequent at 24h (26% compared to 6h (21%) (p<0.01) (Fig. 1B; Fig. S1). These findings indicate that compartment structure destabilizes by 6h and partially normalizes by 24h of reperfusion after ischemia.

**Fig. 1:**
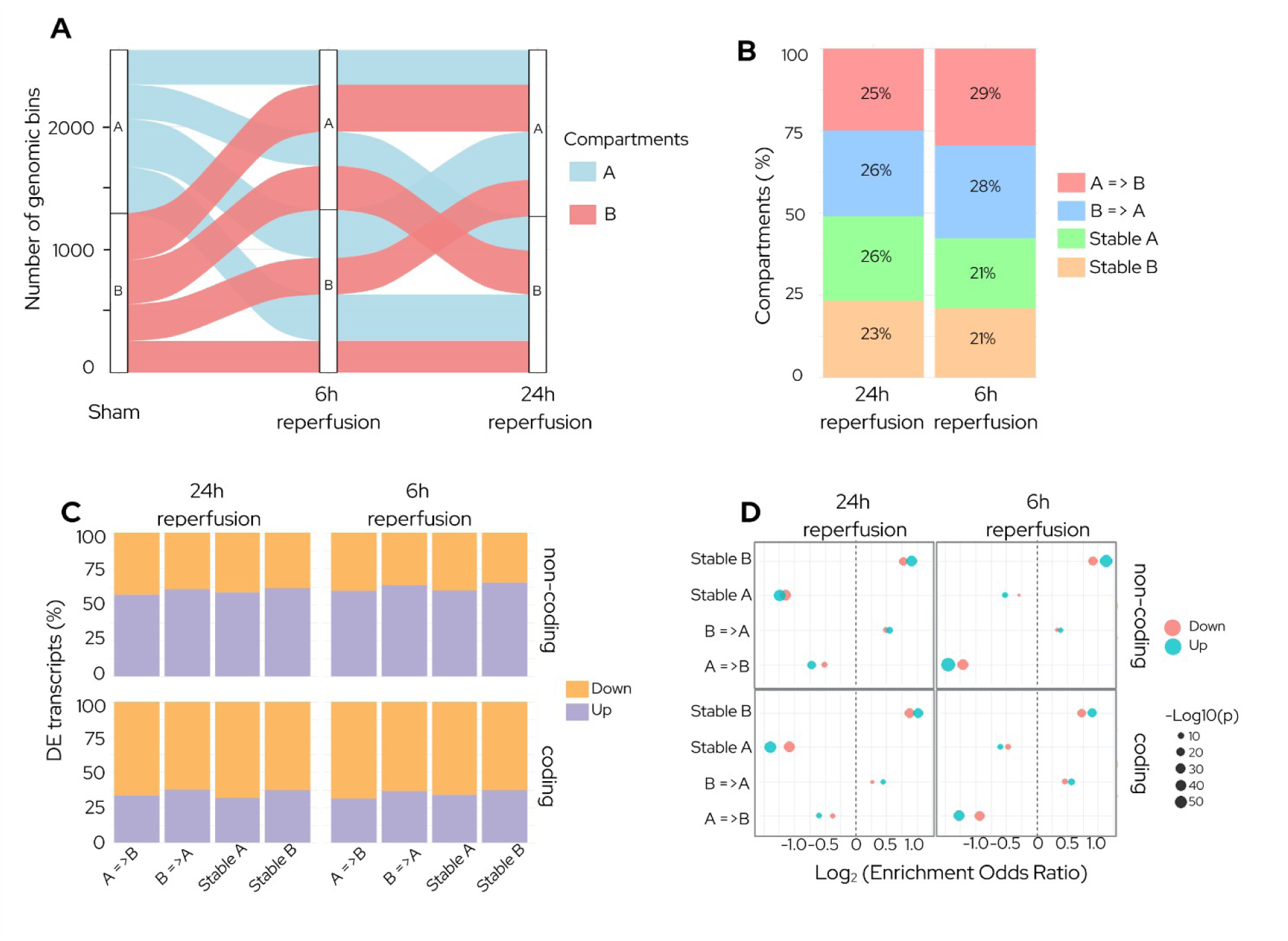
Genome-wide A/B compartment remodeling coordinates early transcriptional responses after stroke. **(A)** Chromatin A/B compartment organization underwent rapid reconfiguration at 6h reperfusion compared to sham, followed by partial recovery at 24h following transient MCAO. **(B)** Proportion of genomic bins classified as A to B, B to A, Stable A, or Stable B at 6h and 24 h reperfusion. Overall compartment switching is higher at 6h reperfusion than at 24h, driven primarily by an increased fraction of A to B transitions. In contrast, the proportion of Stable A compartments increased at 24h reperfusion compared with 6h, indicating a partial recovery toward the pre-stroke compartmental organization pattern. **(C)** Fraction of differentially expressed transcripts within each compartment class at 6h and 24h reperfusion, stratified by transcript type. Noncoding RNAs were predominantly upregulated (>50%) across all compartments, whereas coding transcripts were downregulated (>60%). **(D)** Enrichment of differentially expressed transcripts across compartments, shown as log_2_ enrichment odds ratios for upregulated (teal) and downregulated (salmon) transcripts. Point size reflects –log_10_(p). (n = 3/group).

To determine how transcriptional changes align with the compartments in the ischemic brain, we compared differential expression profiles across compartment classes. Noncoding RNAs exhibited a bias toward upregulation across all compartments (>50%), whereas coding transcripts were predominantly downregulated (>60%) at both reperfusion time points after focal ischemia (Fig. 1C). Specifically, upregulation of noncoding RNAs was strongest in stable B compartments at 6h (adj.p<10⁻^50^) that persisted at 24h (adj.p<10^−30^) of reperfusion. In contrast, coding transcripts showed progressive downregulation from 6h (adj.p < 10⁻^25^) to 24h (adj.p < 10^−50^) reperfusion after focal ischemia (Fig. 1D). Together, these results highlight early noncoding RNA activation in stable B compartments, while compartment switching contributes to subtler, direction-specific modulation of coding transcripts after ischemia.

### TADs are dynamically remodeled in the post-stroke brain

Transient focal ischemia significantly increased the number of TADs (at a resolution of >10 kb) at both 6h and 24h reperfusion compared with sham (p<0.01) (Fig. 2A). While gained TADs dominated at both timepoints (∼75%), lost and shifted TADs were rare (<10%). In the post-stroke brain, ∼77% of the lost TADs resulted from boundary splitting at both reperfusion time points (Fig. 2C). In contrast, most gained TADs formed de novo (>70%), with split-derived TADs being ∼18% at 6h and ∼29% at 24h (Fig. 2C). These findings indicate that stroke drives de novo formation of insulated chromatin domains, independent of pre-existing structures.

**Fig. 2:**
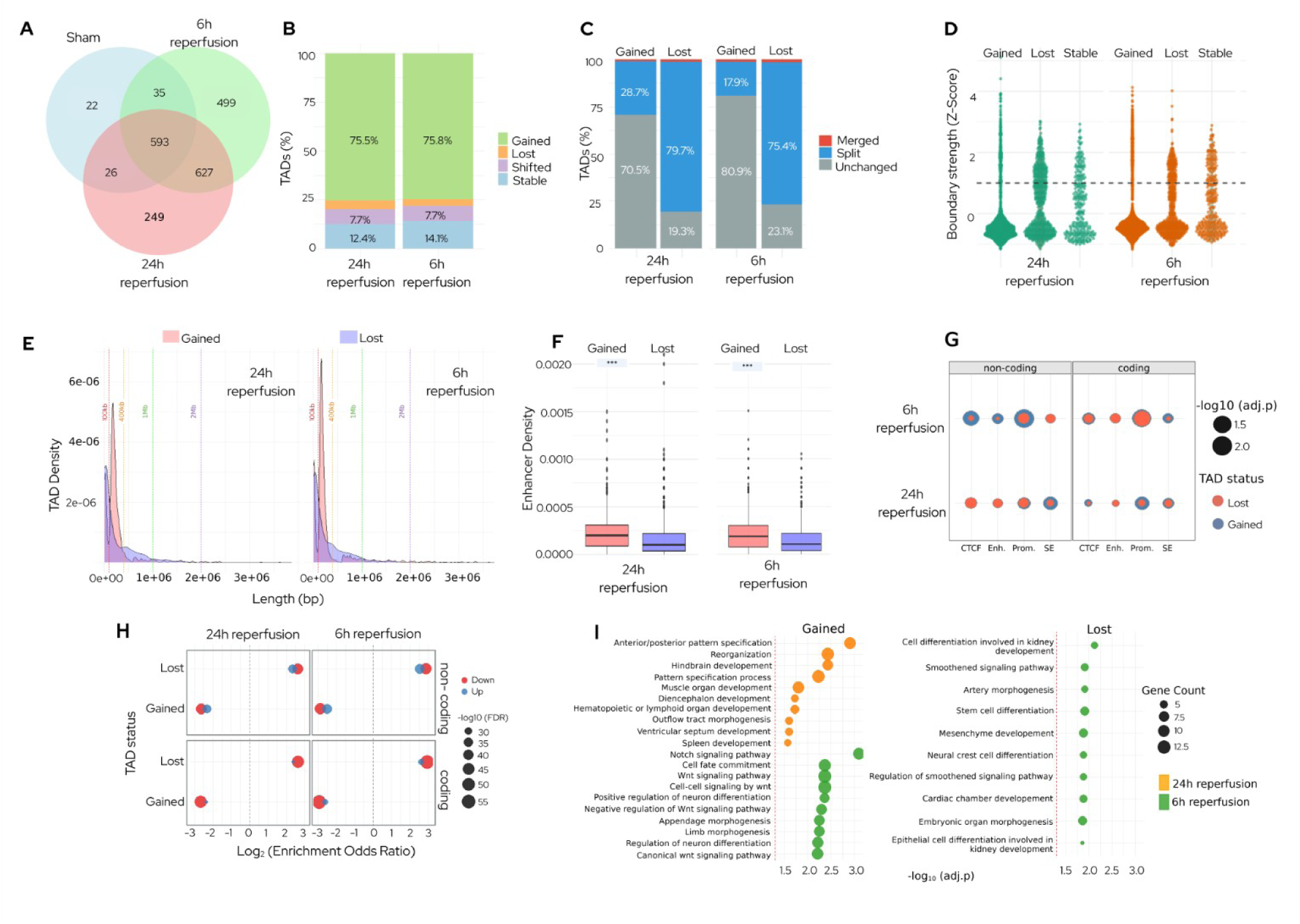
Post-stroke TAD boundaries show dynamic remodeling, accompanied by corresponding changes in stratified functional enhancer landscapes. **(A)** Transient ischemia increased the number of TADs (>10 kb) at both 6h and 24h compared with sham. **(B)** Proportion of TADs gained, lost, or shifted at 6h and 24h reperfusion. ∼75% of TADs are gained, while lost and shifted TADs each were <10%. **(C)** Lost TADs predominantly arose from boundary splitting, while most gained TADs formed de novo after stroke. **(D)** Boundary strength (Z-score) of gained and lost TADs at 6h and 24h reperfusion. Gained TAD boundaries were weaker than lost TADs, with a subset strengthening by 24h (Z-score>2). **(E)** Gained TADs were mostly intermediate in length (100-400 kb), whereas lost TADs were either <100 kb or >400 kb. **(F)** Gained TADs exhibited higher regulatory element density at both reperfusion time points after stroke. **(G)** Association of cis-regulatory elements in gained/lost TADs with noncoding RNA transcripts at 6h. Presence of promoters correlated with a higher number of transcripts. **(H)** Differential transcript regulation in gained and lost TADs. Lost TADs showed dominant downregulation of coding transcripts, whereas gained TADs didn’t show enrichment of differentially expressed transcripts. **(I)** GO analysis of gained and lost TADs at 6h and 24h reperfusion following focal ischemia. Gained TADs showed enrichment in developmental and morphogenetic processes, including Wnt/Notch signaling and neuronal differentiation, with a shift from immune-related functions at 6h and towards tissue patterning at 24h. In contrast, lost TADs were linked to mesenchymal and neural crest differentiation, hedgehog signaling, and organ morphogenesis at both time points.

A subset of the newly formed TAD boundaries strengthened by 24h of reperfusion (boundary Z-score>2) (Fig. 2, Table S3). Additionally, TADs gained after stroke were of intermediate length (100-400kb), whereas TADs lost were either <100KB or >400kb in length (Fig. 2E). Despite reduced boundary strength, gained TADs harbored significantly higher densities of enhancers, promoters, and stroke-specific SEs compared with lost TADs at both reperfusion time points (p<0.01) (Fig. 2F, Fig. S2). Within gained TADs, enhancer and CTCF densities were significantly higher at 24h compared to 6h reperfusion after stroke (p<0.05) (Fig. S3). At 6h reperfusion, the presence of promoters in gained TADs was significantly associated with a higher number of noncoding RNA transcripts (p<0.01) (Fig. 2G). These findings suggest that newly formed TADs, though structurally weak, might be the early hubs of cis-regulatory elements that modulate gene expression in the post-ischemic brain.

TADs lost after stroke showed a dominant transcriptional downregulation, particularly for coding transcripts (adj.p <10⁻^50^ at both reperfusion time points) (Fig. 2H; Fig. S4). In contrast, TADs gained in the post-stroke brain did not show enrichment for differentially expressed transcripts, despite their higher regulatory element density. These results suggest that TAD boundary loss correlates with downregulation of transcripts in the post-stroke brain.

Gene ontology (GO) analysis indicated distinct temporal patterns linked to structural remodeling after stroke (Fig. 2I). Gained TADs enriched for developmental and morphogenetic processes, including Wnt/Notch signaling and neuronal differentiation, and transitioned from immune-related functions at 6h to patterning and tissue-organization pathways at 24h reperfusion. In contrast, lost domains were associated with mesenchymal and neural crest cell differentiation, hedgehog signaling, and organ morphogenesis at both reperfusion time points. These findings demonstrate that ischemia triggers a coordinated 3D genome reorganization that partitions transcriptional programs according to structural and temporal context.

### Chromatin loop remodeling promoted post-ischemic regulatory responses

We detected 16,603 high-confidence reproducible loops at 10 kb resolution in sham which increased significantly (p<0.01) by ∼20% at 6h reperfusion after stroke compared to sham. The number of loops observed at 24h reperfusion was not significantly different from sham. Of these, 4,678 and 2,282 loops were newly formed at 6h and 24h, respectively, reflecting a transient expansion of loops after stroke (Fig. 3A). There was a significant increase in the number of short loops (<200 kb; p<0.05) as well as long loops (≥200 kb; p<0.01) at 6h reperfusion after stroke compared with sham (Fig. 3B and 3C; Fig. S5 and Fig. S6; Table S4). These changes persisted at 24h reperfusion compared with sham for long loops (P<0.01), whereas short loops were not significantly different from sham (Fig. 3B and 3C; Fig. S5 and Fig. S6; Table S4). Across both loop classes and reperfusion time points, more than half of the loops remained stable, while the remainder exhibited distinct gain and loss dynamics (Fig. 3B-C; Fig. S7, Table S5). At 6h reperfusion, ∼29% of long and short loops were gained, while ∼14% were lost, relative to sham (Fig. 3B). By 24h, both gains and losses were ∼18% compared to sham (Fig. 3B).

**Fig. 3:**
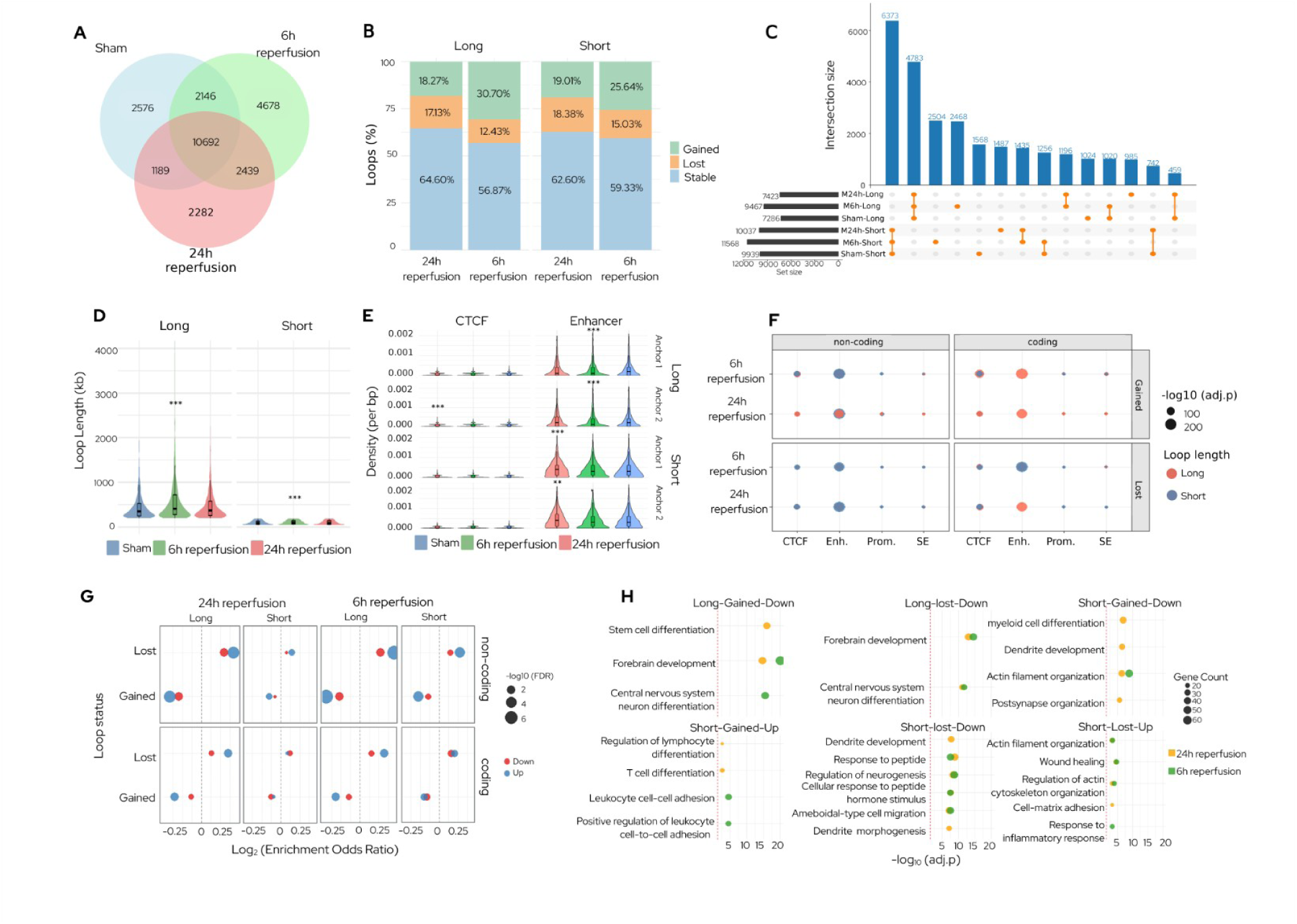
Stroke remodeled chromatin loops to coordinate regulatory responses. **(A)** Number of high-confidence chromatin loops (10 kb resolution) increased transiently at 6h reperfusion compared with sham. **(B)** Numbers of long loops (>200 kb) increased significantly after focal ischemia at both 6h and 24h reperfusion. Whereas number of short loops (<200 kb) increased significantly only at 6h reperfusion. **(C)** Ratios of gained-to-lost short and long loops peaked at 6h reperfusion (short OR=1.65, 95% CI 1.53-1.77; long OR=2.32, 95% CI 2.12-2.52) and normalized by 24h, indicating a reversible burst of looping. More than half of loops remained stable across both reperfusion time points. **(D)** Both short and long loops were significantly elongated at 6h reperfusion following focal ischemia compared with sham. The loop lengths were not significantly different from sham at 24h reperfusion. **(E)** At 6h reperfusion following focal ischemia, enhancer density increased significantly for long loops at both anchor 1 and anchor 2 and for short loops at anchor 2. At 24h reperfusion following focal ischemia, enhancer density remained elevated at both anchors of short loops. CTCF density increased at anchor 2 of long loops at 24h reperfusion following focal ischemia. **(F)** There was a strong enrichment of enhancers for both coding and noncoding RNAs in the gained short loops at 6h reperfusion and gained long loops at 24h reperfusion following stroke. **(G)** Association of loops with differentially expressed transcripts. Lost loops were enriched for upregulated coding and noncoding RNAs at both 6h and 24h reperfusion. **(H)** GO analysis of differentially expressed mRNAs in the gained and lost loops at 6h and 24h reperfusion after focal ischemia. At both 6h and 24h, downregulated mRNAs in long loops (gained and lost) were enriched for forebrain development and neuronal differentiation. Short loops exhibited later enrichment: by 24h, downregulated mRNAs were associated with post-synaptic organization, dendrite development, and actin filament organization. Upregulated mRNAs in gained short loops transitioned from immune cell adhesion at 6h to lymphocyte differentiation at 24h, whereas upregulated short lost loops at 24h were enriched for wound healing, cytoskeletal remodeling, cell adhesion, and inflammatory processes.

Loop length distribution shifted towards a longer span at 6h compared with sham (p<0.01) (Fig. 3D). Gained-to-lost ratios of short and long loops increased significantly at 6h reperfusion compared with sham (short OR=1.65, 95% CI 1.53-1.77; long OR=2.32, 95% CI 2.12-2.52) with partial normalization by 24h reperfusion (short OR=1.00, 95% CI 0.93-1.08; long OR=1.00, 95% CI 0.92-1.09) (Fig. 3D). This suggests that stroke triggers a rapid widespread, but transient loop remodeling, characterized by early gains followed by partial normalization of 3D genome architecture by 24h reperfusion.

Following stroke, enhancer density increased significantly at 6h reperfusion for anchor 1 and anchor 2 for long loops (p<0.01 for both) and anchor 2 for short loops (p<0.01) and remained elevated at 24h reperfusion for both anchor 1 and anchor 2 for short loops (p<0.01 for both) compared with sham (Fig. 3E; Fig. S8). CTCF density increased at anchor 2 of long loops (p<0.01), while promoter density remained unchanged at 24h reperfusion compared with sham (Fig. 3E; Fig. S8). At both reperfusion time points after ischemia, regulatory enrichment association with the transcriptome profile of coding and noncoding RNAs followed a consistent hierarchy, with significant enrichment for enhancers > CTCF > promoters ∼ SEs (Fig. 3F, Fig. S8).

Enhancer annotations were significantly enriched in gained loops for both coding and noncoding RNAs with high enrichment in short loops at 6h reperfusion (adj.p<10⁻^200^) and for coding RNAs in long loops at 24h reperfusion (adj.p<10⁻^200^) (Fig. 3F). At 6h reperfusion, transcripts located in gained long loops were strongly associated with CTCF-binding regions (adj.p<10^−200^), whereas associations with promoters were comparatively modest across all loop classes (adj.p<10^−5^) (Fig. 3F). Overall, Stroke promoted changes in enhancer occupancy at loop anchors, along with CTCF gains and selective super-enhancer associations at 6h reperfusion. Enhancer-containing loops exhibited the strongest link to transcript-level differential expression, highlighting enhancer-loop associations as the principal regulatory signature of post-ischemic loop dynamics. Despite their regulatory enrichment, gained loops were less frequently associated with differential gene expression compared to lost loops in the post-stroke brain. Following transient MCAO, upregulated noncoding RNAs were enriched in lost loops (6h: OR=1.37, adj.p<10^−5^; 24h: OR=1.34, adj.p<10^−5^) and depleted in gained loops (6h: OR=0.73, adj.p<10^−5^; 24h: OR=0.75, adj.p<10^−5^) (Fig. 3G). Thus, loop dissolution marks loci of active transcriptional regulation following stroke.

GO analysis of differentially expressed mRNAs within gained and lost loops showed time- and loop-type-specific transcriptional reorganization in processes relevant to stroke pathophysiology (Fig. 3H; Fig. S9). Downregulated mRNAs in gained and lost long loops at both 6h and 24h reperfusion were enriched for forebrain development and neuron differentiation, indicating suppression of neurodevelopmental pathways. In gained short loops at 24h, downregulated mRNAs were associated with post-synaptic organization, dendrite development, actin filament organization, and myeloid lineage differentiation. Concurrently, upregulated mRNAs in gained short loops shifted from immune cell adhesion at 6h to lymphocyte differentiation by 24h reperfusion. Lost short loops displayed a parallel pattern in which downregulated mRNAs were enriched for neuronal structure and morphogenesis, while upregulated mRNAs were associated with wound healing, actin cytoskeleton remodeling, cell adhesion, and inflammatory responses at 24h reperfusion. Together, these findings suggest that loop remodeling coordinates a transcriptional transition from neurodevelopmental and synaptic processes to immune activation and tissue repair, with early changes in long loops at 6h and a shift in short loops by 24h reperfusion after stroke.

### Integrated Multi-scale Remodeling Links Structure to Gene Regulation

Across TAD contexts and time points, the proportion of gained versus lost loops varied (Fig. 4A). Additionally, loop remodeling after stroke was structured by TAD context and loop length. At 6h reperfusion, gained short loops were predominantly located outside TADs (∼57%), with additional contributions from inside-TAD loops (∼30%) and one-anchor loops (∼10%), whereas loops bridging distinct TADs were rare (∼3%). Lost short loops displayed a similar distribution, with ∼53% outside TADs, ∼37% inside TADs, ∼9% one-anchor, and ∼0.6% inter-TAD loops. Long loops gained or lost at 6h were primarily outside TADs (∼39%) or one-anchor (∼31%), followed by inside-TAD (∼29%) and inter- TAD loops (∼1.5%) (Fig. S10). By 24h reperfusion, the proportional landscape of loops was largely stable. Gained short loops remained enriched outside TADs (∼58%), followed by inside-TADs (∼31%), one-anchor (∼9%), and inter-TAD loops (∼2%) (Fig. S10). Lost short loops mirrored the 6h profile. For long loops, both gains and losses were dominated by outside-TADs (∼38%) and one-anchor loops (∼31%), with 29% inside TADs and 2% bridging TADs (Fig. S10). Overall, across both time points and loop classes, gain and loss events were enriched outside TADs, whereas inter-TAD loops were a minor fraction (∼2%). These results indicate that stroke-driven loop remodeling predominantly occurs within existing TAD boundaries rather than formation of de novo inter-domain bridges. Compositionally, outside-TAD loops contributed to the largest share of loop gains at both reperfusion time points after focal ischemia. However, gains within TADs also represent a substantial fraction, indicating that domain-contained loop remodeling events are common (Fig. 4A; Fig. S10). These data suggest that post-stroke loop dynamics are structured by TAD context, with *de novo* interactions arising both within and beyond domain boundaries.

**Fig. 4:**
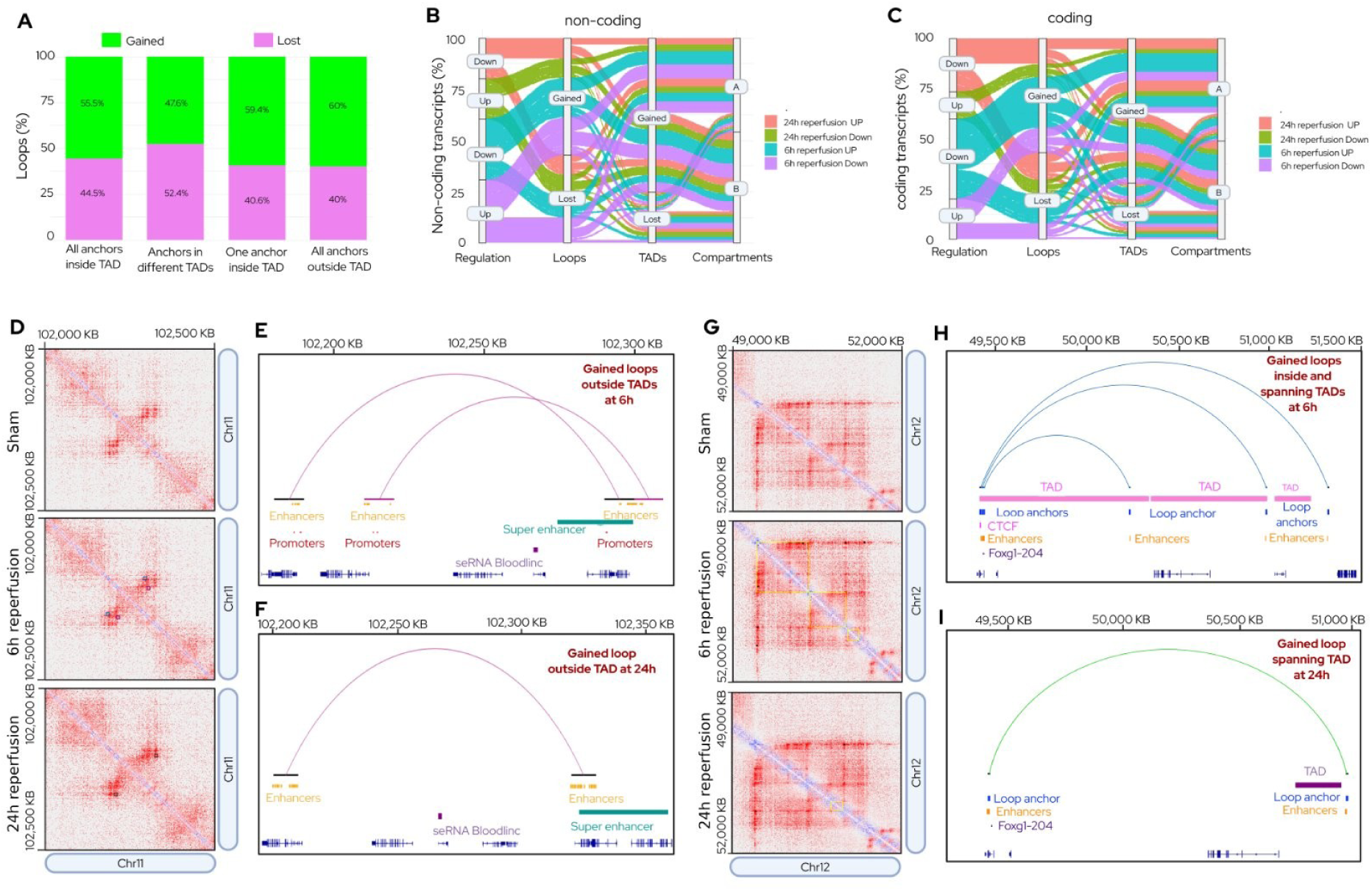
Multi-scale chromatin remodeling links loop/TAD dynamics to transcriptional regulation in the post-stroke brain. **(A)** Proportion of loops in different genomic contexts (inside TAD, outside TAD, one-anchor within a TAD, bridging two TADs). Percentages of gained and lost loops are indicated for each category. Outside-TAD loops contributed the largest fraction of gains. Short (<200 kb) and long (≥200 kb) loops were combined. **(B–C)** Sankey diagrams depicting the hierarchical architecture of transcript regulation through loops to TADs to compartments for noncoding RNAs (B) and coding RNAs (C). Percentages along paths indicate the fraction of transcripts following each regulatory route. At both 6h and 24h reperfusion following transient MCAO, upregulated transcripts were most frequently associated with gained loops flowing into gained TADs as well as lost loops flowing into gained TADs. Downregulated transcripts also showed a similar distribution. **(D)** Heatmaps for sham, 6h, and 24h reperfusion times with annotated TADs and loops in the Bloodlinc locus (chr11:102–102.5 Mb). **(E)** At 6h reperfusion after focal ischemia, Bloodlinc locus was observed to be located within two gained loops outside TADs spanning enhancers, promoters, and a stroke-specific super-enhancer. **(F)** By 24h, Bloodlinc locus was encompassed by a single gained loop with a set of enhancers and a super-enhancer. **(G)** Heatmaps for sham, 6h, and 24h reperfusion times with annotated TADs and loops for Foxg1 locus (chr12: ∼49–52 Mb). **(H)** At 6h (Foxg1 locus), three gained loops spanned adjacent TADs of which one anchored by CTCF. **(I)** By 24h, Foxg1 locus loops consolidated into a single TAD-crossing loop with fewer enhancers and no distal CTCF, illustrating loop/TAD reconfiguration while maintaining transcriptional activity.

Sankey plots tracing transcript regulation through the hierarchical architecture of loops to TADs to compartments revealed diverse regulatory flows for both noncoding RNAs (Fig. 4B) and coding RNAs (Fig. 4C). Many upregulated transcripts resided in gained loops within gained TADs that were retained in the A compartment. Across reperfusion groups, transcript regulation was dominated by TAD gains. At 6h reperfusion, upregulated transcripts were most frequently associated with gained loops flowing into gained TADs (∼46%), with an additional ∼33% corresponding to lost loops in gained TADs (Fig. 4B-C, Table. S6). Downregulated transcripts exhibited a similar pattern, with ∼45% in gained loops and ∼29% in lost loops within gained TADs. By 24h reperfusion, the distribution of up- and downregulated transcripts approached a more balanced state, with ∼38% of upregulated and ∼37% of downregulated transcripts in gained loops, and ∼38% of upregulated and ∼40% of downregulated transcripts in lost loops flowing into gained TADs, reflecting a shift towards equilibrium in loop-mediated transcript localization (Fig. 4B-C, Table. S6).

When analyzed in the full hierarchical context, including compartment identity, upregulated transcripts were frequently localized in gained loops to gained TADs (∼44%), followed by lost loops to gained TADs (∼34%), gained loops to lost TADs (∼14%), and lost loops to lost TADs (∼11%). Downregulated transcripts displayed a similar distribution, with a gain-bias at 6h that diminished by 24h, while TAD-level remodeling continued to dominate transcript localization. These results indicate that transcript regulation after stroke reflects coordinated multi-scale chromatin reorganization, with TAD-level changes exerting a more persistent influence than loop turnover alone.

Locus exemplars (Bloodlinc and Foxg1) illustrated time-resolved gene regulation after stroke. Bloodlinc is a SE-driven long noncoding RNAs upregulated significantly (by ∼3.7-fold) at both 6h and 24h reperfusion compared with sham (Table S7). At 6h, the Bloodlinc locus was between the anchors of 2 gained loops near a stroke-specific SE which reorganized into a dominant loop enclosing Bloodlinc and repositioning it nearby a second SE by 24h reperfusion (Fig. 4D–F). In both cases, the anchors overlap SEs without intervening TAD boundaries (i.e., outside TAD), permitting sustained enhancer-promoter communication. Foxg1, a transcription factor that modulates neurogenesis was upregulated by ∼27-fold at 6h and by ∼21-fold at 24h reperfusion compared with sham (Table S7). The Foxg1 locus was near 3 gained loops spanning adjacent TADs, one anchored by a CTCF site at 6h reperfusion. By 24h reperfusion, this configuration condensed into a single loop crossing a TAD boundary, with fewer enhancers and loss of the distal CTCF (Fig. 4G–I). This locus exemplifies how loop/TAD reorganization might lead to reduced transcription after stroke.

## Discussion

Ischemic stroke initiates a rapid and hierarchical remodeling of chromatin architecture, orchestrating temporally resolved transcriptional reprogramming. By integrating high-resolution Hi-C data with transcriptomics data, we presently show that genome topology (including compartments, TADs and loops) undergoes a coordinated, scale-dependent reorganization after stroke. These structural transitions are not merely reflective of transcriptional states, but actively shape gene responsiveness, particularly of noncoding RNAs. Stroke induced a selective loss and gain of chromatin domains across hierarchical scales. At the loop level, gained loops were not uniformly associated with transcriptional activation in the post-stroke brain. At the TAD level, gained TADs serve as a principal scaffold through which regulatory flows are routed. Integrating across scales, we observed that gained loops nested within gained TADs might preferentially support transcriptional regulation which underscores the importance of architectural context in determining loop functionality. Together, these findings establish time-dependent genome remodeling as a critical layer of transcriptional regulation in the post-stroke brain and provide a framework for decoding stimulus-dependent chromatin plasticity.

At the megascale, we observed a pronounced A to B compartment shift at 6h reperfusion, that partially stabilized by 24h reperfusion after stroke. This transient A to B compartment switching might shape the early post-ischemic transcriptional responses. We observed that stable B compartments, considered repressive, were enriched for upregulated transcripts, particularly noncoding RNAs, at both reperfusion timepoints after stroke compared with sham. This challenges the conventional model of B compartments as silencing and supports flexible, stimulus-responsive gene regulation by a more dynamic B domains.^29,30^

While B compartments are known to be associated with lower transcriptional activity, subsets of loci within them retain expression competence. For example, in B cell lymphoma, compartment switching was shown to affect only a subset of genes, and B compartments harbored genes with relatively high expression levels.^31^ Presently observed switching of transcripts from A to B compartments suggests a repressive influence, potentially mediated by lamina-association or chromatin compaction in the post-stroke brain. This is consistent with previous studies that showed that compartmental status contributes to, but does not fully determine, cell type-specific transcriptional programs.^32,33^ In a nutshell, our results suggest that B compartments might function as dynamic hubs for stimulus-responsive noncoding RNA activity in the post-ischemic brain.

At the mesoscale, TAD boundaries are highly conserved across mammalian species.^34^ Following stroke, we observed a marked increase in the number of TADs, predominantly due to the emergence of novel domains. Similar boundary rearrangements have been documented during development and in various disease states.^30,32,35–37^ Moreover, a recent study reported structural alterations in TADs in the peri-infarct cortex of mice subjected to focal ischemia.^27^ While a subset of newly formed TADs displayed stronger boundaries at 24h reperfusion, most exhibited weaker boundary strength. Previous studies also showed that nascent TAD boundaries, such as those formed during neural differentiation, mature gradually over time. For example, new TAD boundaries in developing neural tissue form progressively and exhibit increased insulation as CTCF occupancy increases.^35,36^ However, the present study did not observe elevated CTCF motif density in gained TADs. As our analysis was based solely on motif presence, and not binding activity or cohesin co-occupancy, we can’t determine whether these domains are functionally insulated by CTCF or merely represent potential sites of insulation. Many gained TADs represent immature structural units lacking sufficient insulation to support stable enhancer-promoter interactions. Despite this, gained TADs were significantly enriched for enhancers, promoters, and stroke-specific SEs in our dataset. Yet paradoxically, they were not enriched for differentially expressed genes. This apparent disconnect between regulatory element density and transcriptional output may reflect nonlinear enhancer-promoter dynamics. Recent studies showed that promoter activation depends on a bistable regulation in which transient enhancer contact may not immediately trigger gene expression.^38^ Hence, the weak insulation of gained TADs may limit sustained enhancer-promoter communication after stroke, rendering these regions transcriptionally inert.^38,39^ In contrast, lost TADs were enriched for downregulated transcripts, particularly coding genes, suggesting a repressive shift following domain loss during the acute reperfusion phase after stroke. This indicates that TAD boundaries might act as scaffolds to facilitate selective enhancer-promoter communication. Disruption of these boundaries might impair regulatory specificity and results in transcriptional repression in the post-stroke brain, even when contact frequencies remain modestly altered.^38^ This was supported by the observation that deletion of TAD boundaries can lead to widespread downregulation of resident genes in mouse embryos.^40,41^ Our study indicates that post-stroke domain loss preferentially affects transcription of coding genes, likely through compromised enhancer targeting.

We observed that chromatin architecture rapidly reconfigured after stroke, with a substantial increase in loop number and length that partially stabilized by 24h reperfusion. This remodeling resembles stimulus-induced loop dynamics reported in other biological systems such as neural differentiation, and skeletal muscle lineage progression.^35,42–44^ Importantly, our results highlight a functional asymmetry in loop dynamics; gained loops are not uniformly linked to transcriptional activation, whereas lost loops frequently coincide with upregulated transcripts, particularly noncoding RNAs. In the post-stroke brain, despite being enriched for enhancers and promoters, gained loops were not enriched for transcriptional changes. Previous studies also demonstrated that during neural differentiation many gained loops are not immediately linked to gene expression, suggesting that architectural reorganization can occur independently of transcription and may serve as prime loci for later activation of gene expression.^35,45^ In contrast, a study showed that gene expression exhibits a directional bias where gained loops tend to associate with upregulation of genes positioned outside the loop center, whereas lost loops show higher expression of genes located within loop boundaries.^43^ This model could explain our observation that noncoding RNAs are preferentially upregulated in lost loops. In support, we observed that lost loops were strongly associated with transcriptional changes, particularly noncoding RNAs. Although direct evidence linking loop dissolution specifically to noncoding RNA upregulation is limited, a previous study showed that loop loss can increase gene expression.^46^ In embryonic stem cells, loop disruption was reported to enhance the expression of genes.^47^ Moreover, noncoding RNAs are known to exploit architectural remodeling as a mechanism for transcriptional activation.^48^ Thus, noncoding RNA upregulation might actively reshape chromatin structure and genome architecture after stroke.

The regulatory impact of loop dynamics is dictated by the broader multi-scale chromatin context. The transcriptional consequences of loop remodeling were amplified when nested within permissive domains. For instance, loops within gained TADs had significantly higher odds of overlapping with differentially expressed transcripts after stroke. Sankey flow analysis confirmed that gained TADs provide the dominant framework for routing transcriptional changes, while compartment identity adds further specificity, with Stable B and B to A switching enriched for upregulated noncoding RNAs after stroke. Thus, loop remodeling functions in coordination with domain architecture to create permissive regulatory environments in the post-stroke brain.^35,42^ Gene exemplars illustrate these multi-scale principles. The noncoding RNA *Bloodlinc*, positioned outside TAD boundaries, but flanked by gained loops, anchored at stroke-specific SEs and showed sustained upregulation in the post-stroke brain. This indicates that the stabilization of enhancer-promoter communication after stroke might be independent of TAD remodeling. Conversely, *Foxg1* is one of the highly expressed genes despite extensive loop rewiring and TAD boundary crossing, illustrating that transcriptional programs can persist even when architectural configurations shift. Together, these genes emphasize that loop remodeling after stroke operates within a hierarchical chromatin framework, where regulatory outcomes depend on loop status, integration with TADs and compartment context.

Bulk Hi-C and microarray data lack cell-type resolution, which is a limitation of the present study. However, the global view reflects the coordinated response of neurons, glia, and infiltrating immune cells to ischemia. Our findings thus provide a framework to build future single-cell and spatial Hi-C studies. Notably, features such as B-compartment enrichment of noncoding RNAs and loop/TAD-dependent immune activation show tissue-specific changes that can’t be observed with single-cell methods. In the present studies, CTCF, enhancer, and promoter annotations were based on experimentally defined mouse datasets, that are not stroke-specific. In contrast, SEs were mapped directly from peri-infarct cortex and restricted to stroke-unique regions at 24h reperfusion after stroke. Additional integration of cohesin occupancy and chromatin accessibility will show the insulation strength and enhancer competence. Despite these considerations, our study established a genome-wide chromatin plasticity in stroke and points to nuclear architecture as an untapped therapeutic axis for transcriptional recovery.

In summary, the multi-scale mapping of the spatiotemporal chromatin architecture of the present study indicates that genome reorganizes during the acute phase after ischemic stroke. We showed that transcriptional reprogramming is not a global wave of activation, but a hierarchical process in which shift of compartments, loss of TADs and loop remodeling act in concert to selectively coordinate gene expression. Present studies also indicate that injured brain responds through structured architectural funneling of regulatory signals than chaotic genomic disruption.

## Supporting information

Statistics Tables

R packages versions

## Acknowledgments

We thank the University of Wisconsin-Madison Biotechnology Center Sequencing Facility for performing DNA fragmentation of Hi-C samples and sequencing. We thank the University of Wisconsin-Madison Biotechnology Center Bioinformatics Core for quality control of sequencing data and for carrying out Juicer-based analyses, including eigenvector decomposition, Arrowhead, and HiCCUPS.

