## Supplementary Figures for "Hierarchical Chromatin Rewiring Orchestrates Early Transcriptional Regulation in Mouse Cerebral Cortex Following Focal Ischemia"

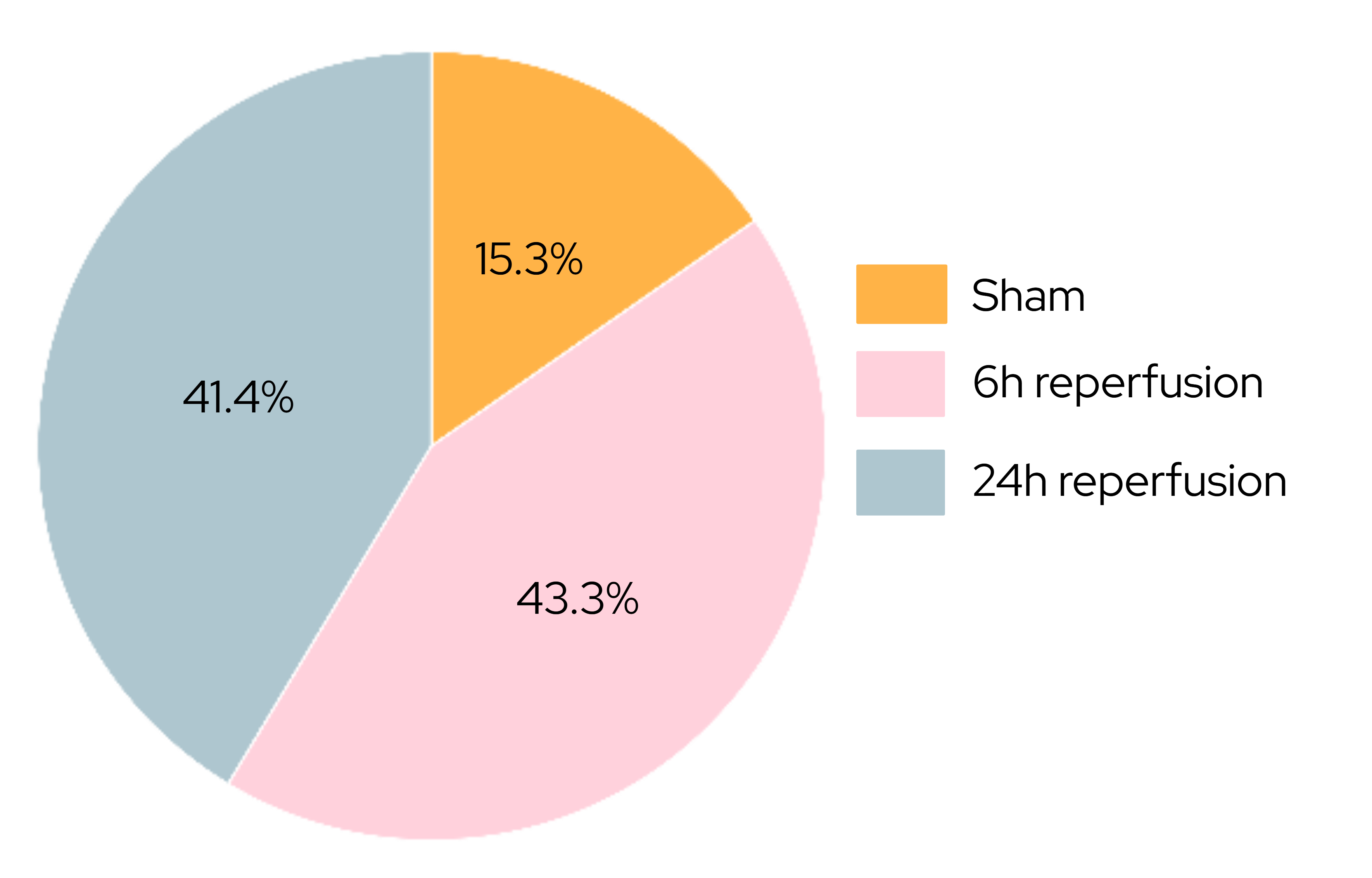

**Figure S1.** Percent distribution of reproducible TADs across experimental conditions. The proportion of reproducible topologically associating domains (TADs) is shown for Sham, 6 h, and 24 h reperfusion following transient MCAO.

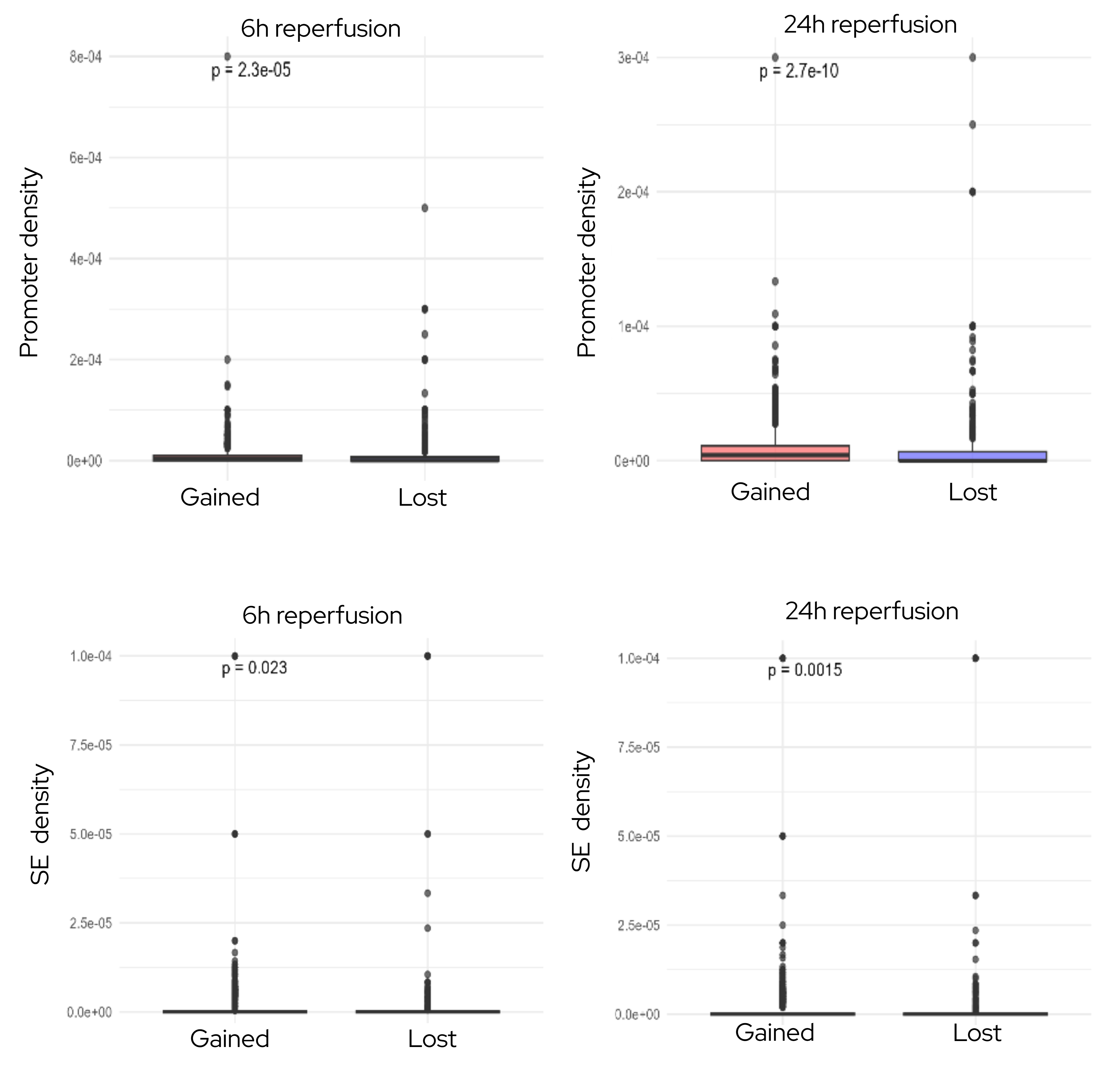

**Figure S2**. Promoter and super-enhancer (SE) densities in gained versus lost TADs after stroke. Promoter and super-enhancer densities were compared between gained and lost TADs at 6 h and 24 h reperfusion following transient MCAO. Statistical significance was assessed using the Mann–Whitney U test. Only differences with p < 0.05 are shown.

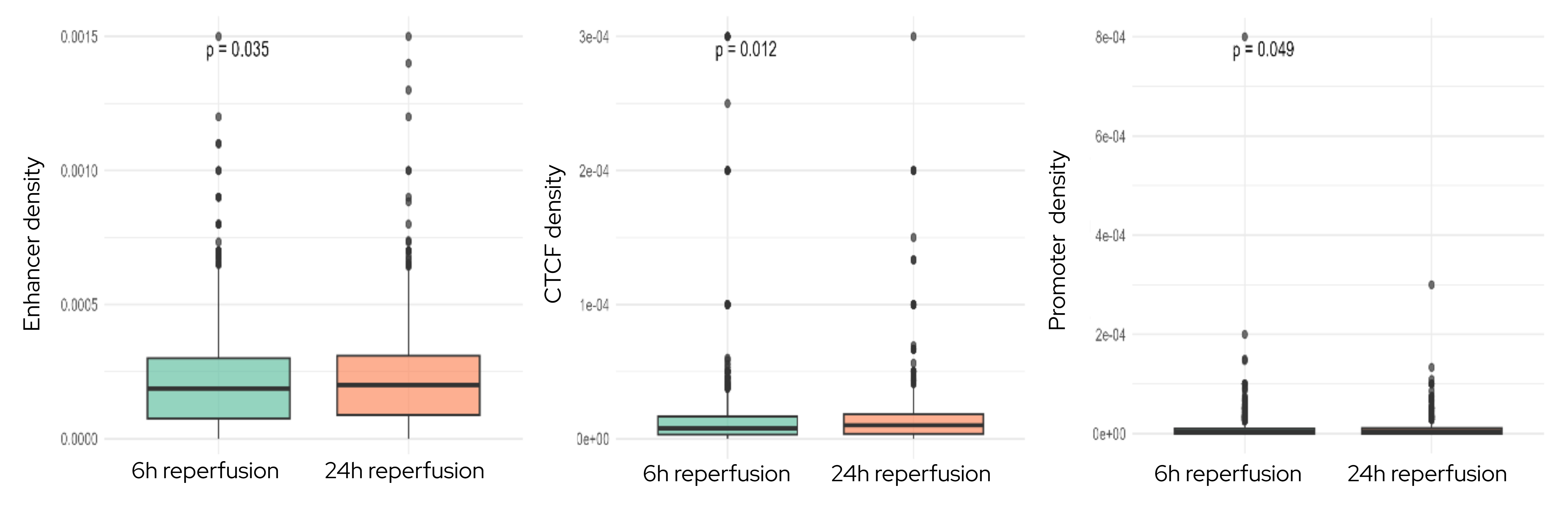

**Figure S3.** Differences in enhancer, CTCF, and promoter densities between gained TADs at 6 h and 24 h reperfusion. Enhancer, CTCF, and promoter densities were compared between gained TADs at 6 h and 24 h reperfusion following transient MCAO. Statistical significance was assessed using the Mann–Whitney U test. Only differences with p < 0.05 are shown.\

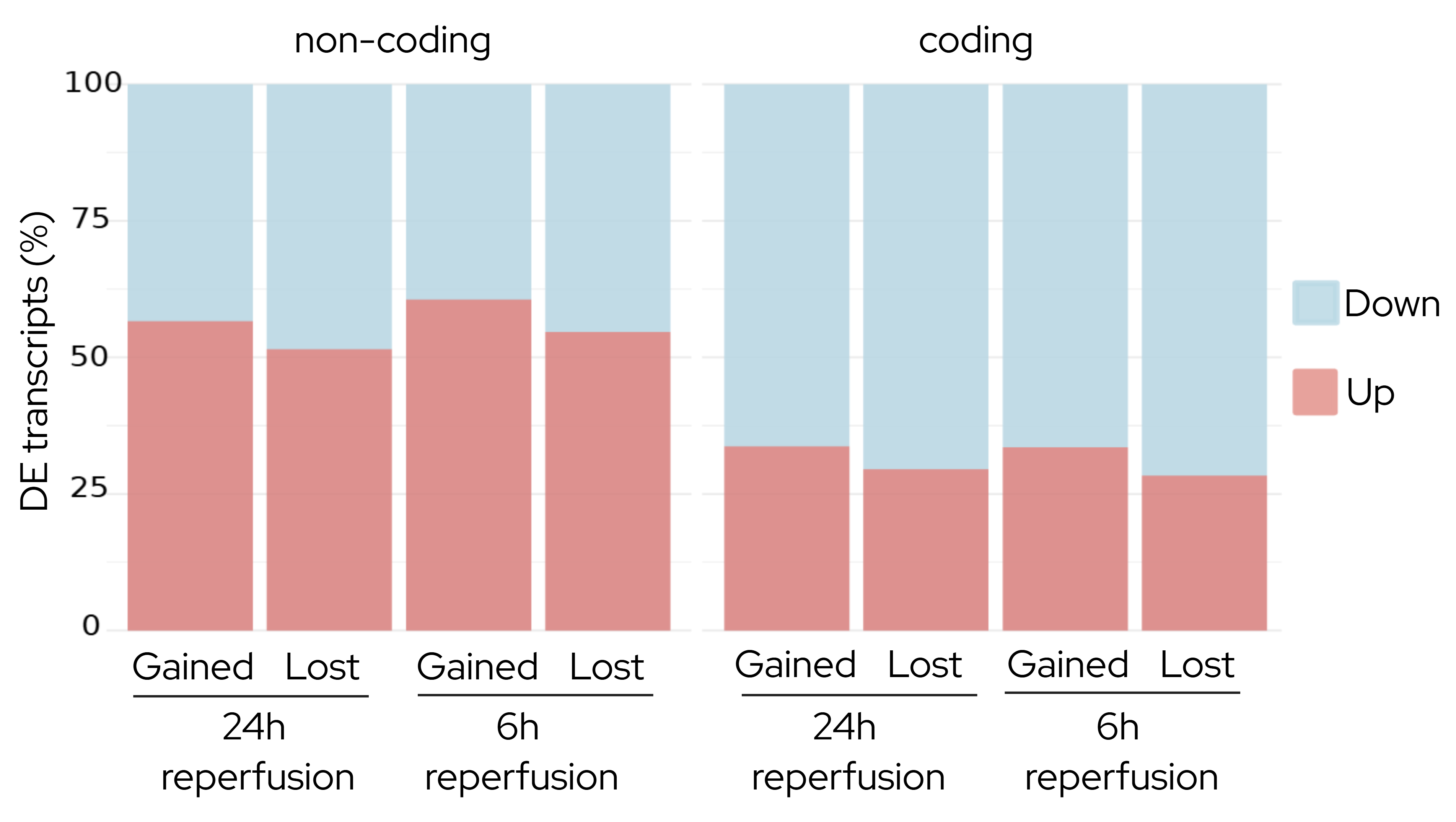

**Figure S4.** Percent distribution of differentially expressed transcripts across TAD categories at 6 h and 24 h reperfusion. The proportion of upregulated and downregulated transcripts is shown for gained and lost TADs at 6 h and 24 h reperfusion following transient MCAO, stratified by protein-coding and non-coding transcripts.

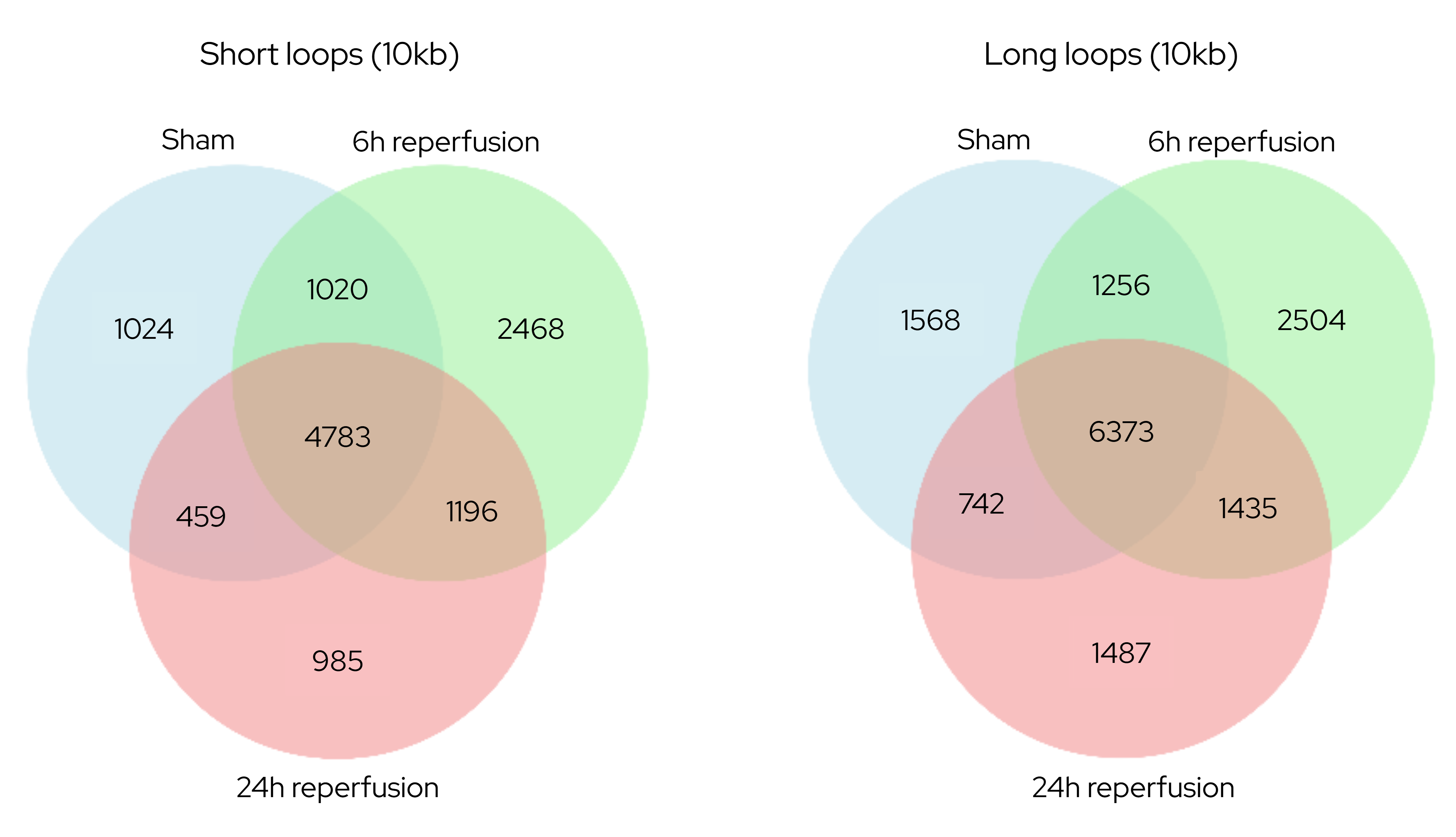

**Figure S5.** Distribution of reproducible short and long chromatin loops across experimental conditions. Venn diagrams show the distribution of reproducible short (<200 kb) and long (≥200 kb) chromatin loops across Sham, 6 h, and 24 h reperfusion following transient MCAO, identified at 10 kb resolution.

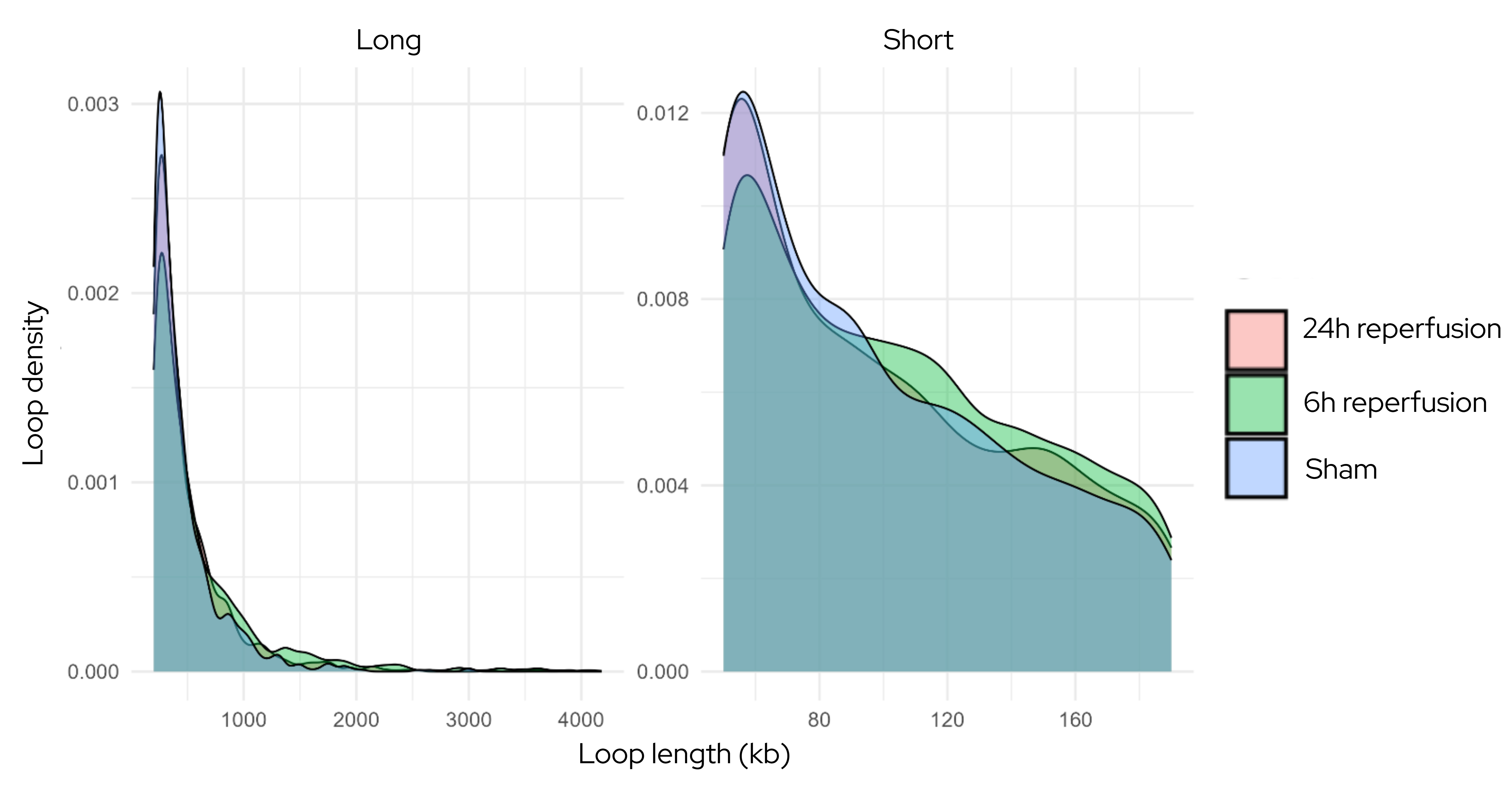

**Figure S6.** Loop length distributions across experimental groups. Loop length density distributions are shown for Sham, 6 h, and 24 h reperfusion following transient MCAO, stratified by short (<200 kb) and long (≥200 kb) chromatin loops.

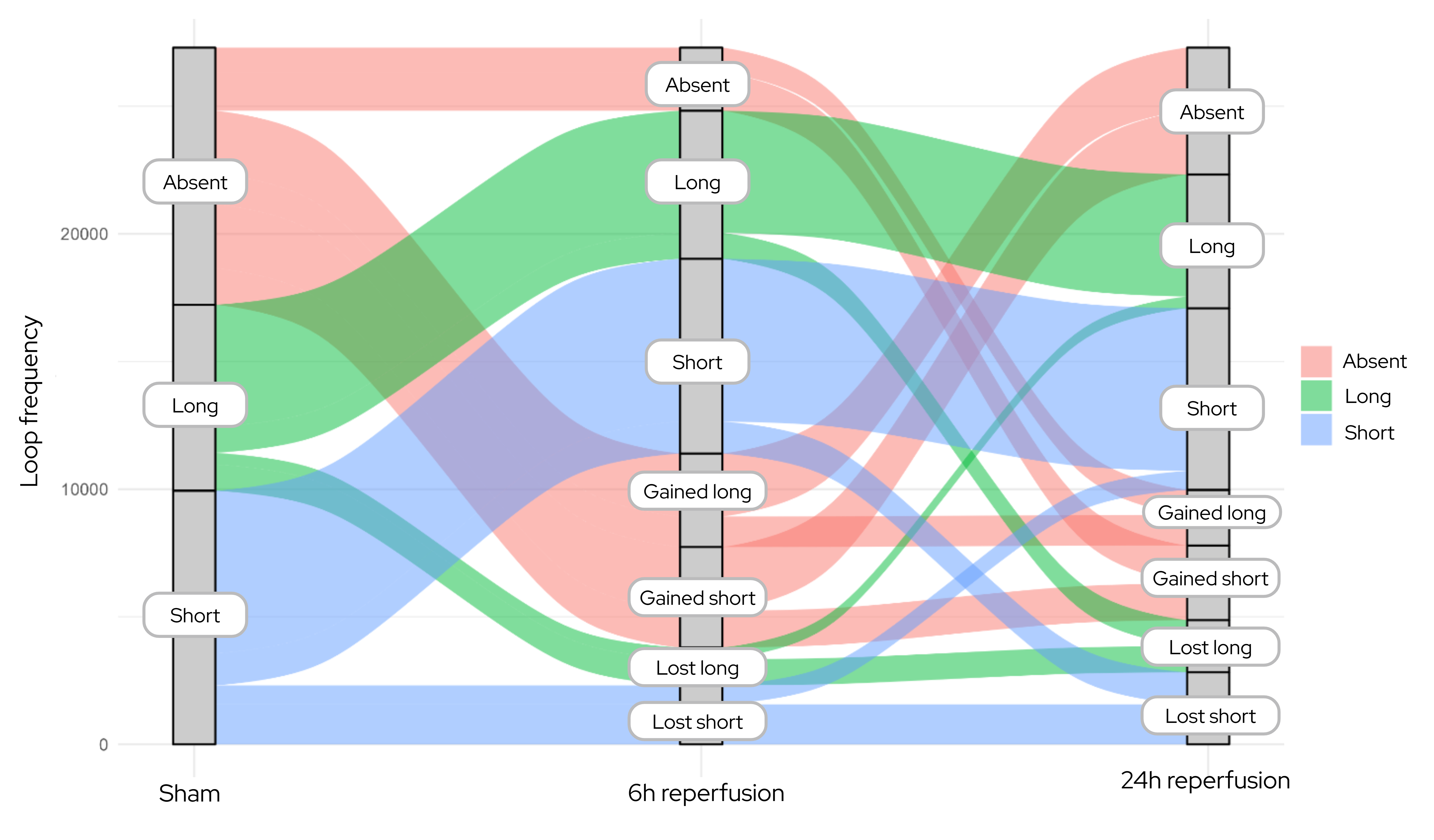

**Figure S7**. Sankey plot illustrating transitions of short and long chromatin loops between experimental groups. The Sankey plot shows the flow of short (<200 kb) and long (≥200 kb) loops across Sham, 6 h, and 24 h reperfusion following transient MCAO. Loops absent in a group are labeled “Absent.” Loops gained at 6 h or 24 h but absent in Sham are labeled “Gained,” whereas loops present in Sham but absent at 6 h or 24 h are labeled “Lost.”

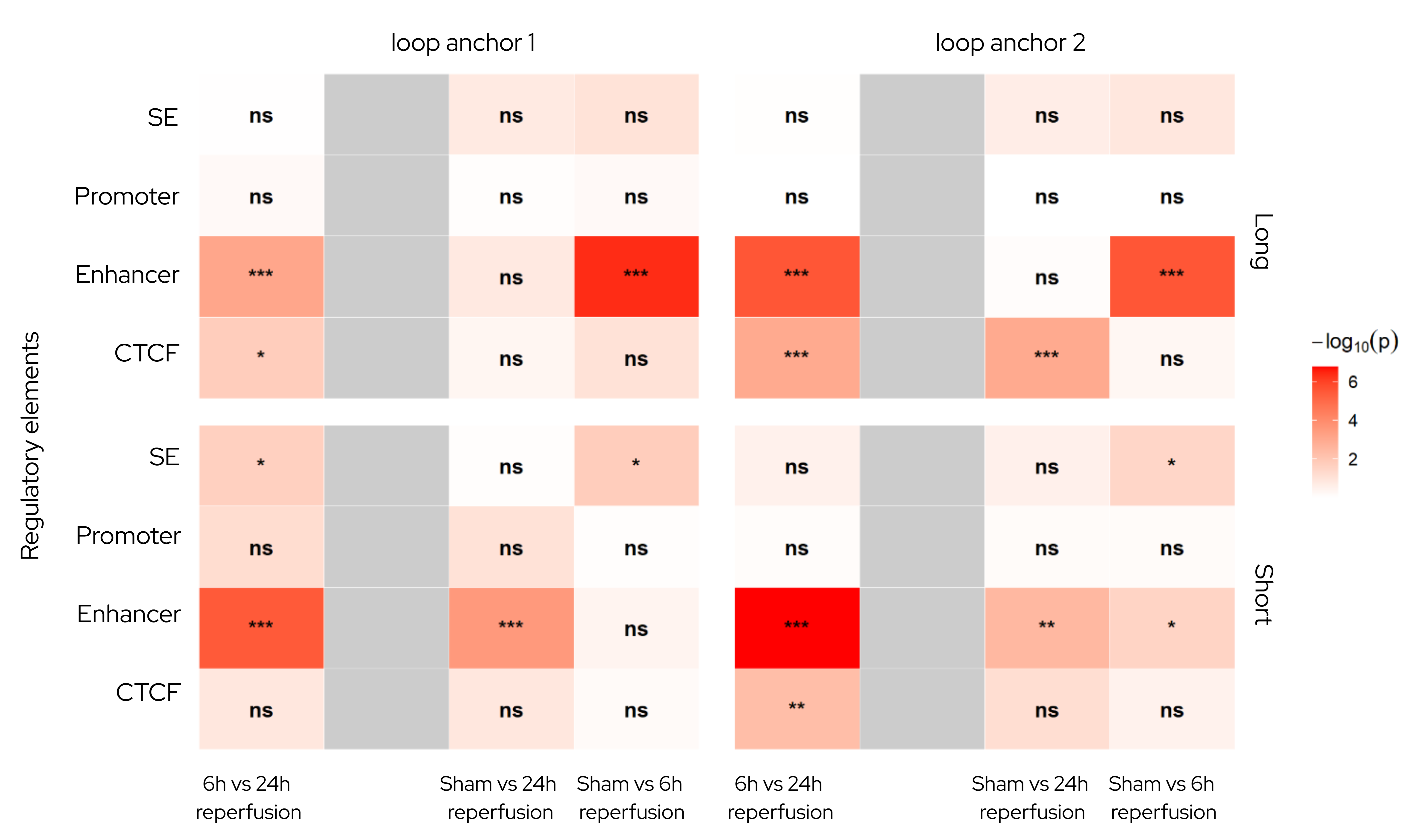

**Figure S8.** Densities of cis-regulatory elements at loop anchors for short and long loops. Heatmap shows densities of super-enhancers (SE), promoters, enhancers, and CTCF sites at both anchors of short (<200 kb) and long (≥200 kb) chromatin loops. Comparisons between experimental groups (Sham, 6 h, 24 h reperfusion following transient MCAO) were assessed using pairwise Mann–Whitney U tests (Wilcoxon rank-sum test equivalent). Color intensity represents –log₁₀(p), with non-significant comparisons labeled “ns.”

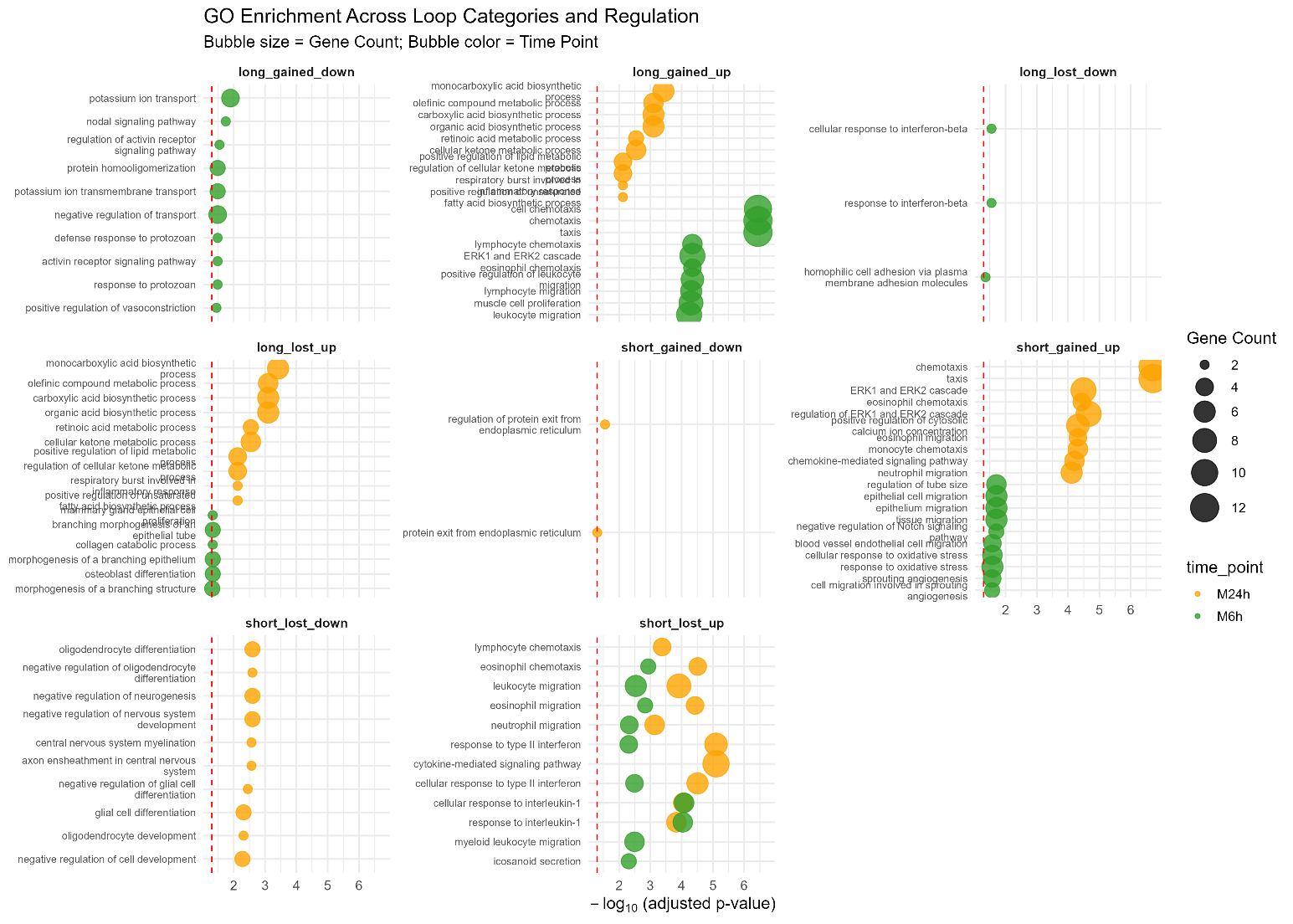

**Figure S9.** Gene Ontology (GO) enrichments for differentially expressed mRNAs by loop category. GO enrichment analysis was performed separately for short (<200 kb) and long (≥200 kb) loops, stratified by experimental group (Sham, 6 h, 24 h reperfusion) and direction of regulation (upregulated or downregulated). The top 20 enriched GO terms for each category are shown.

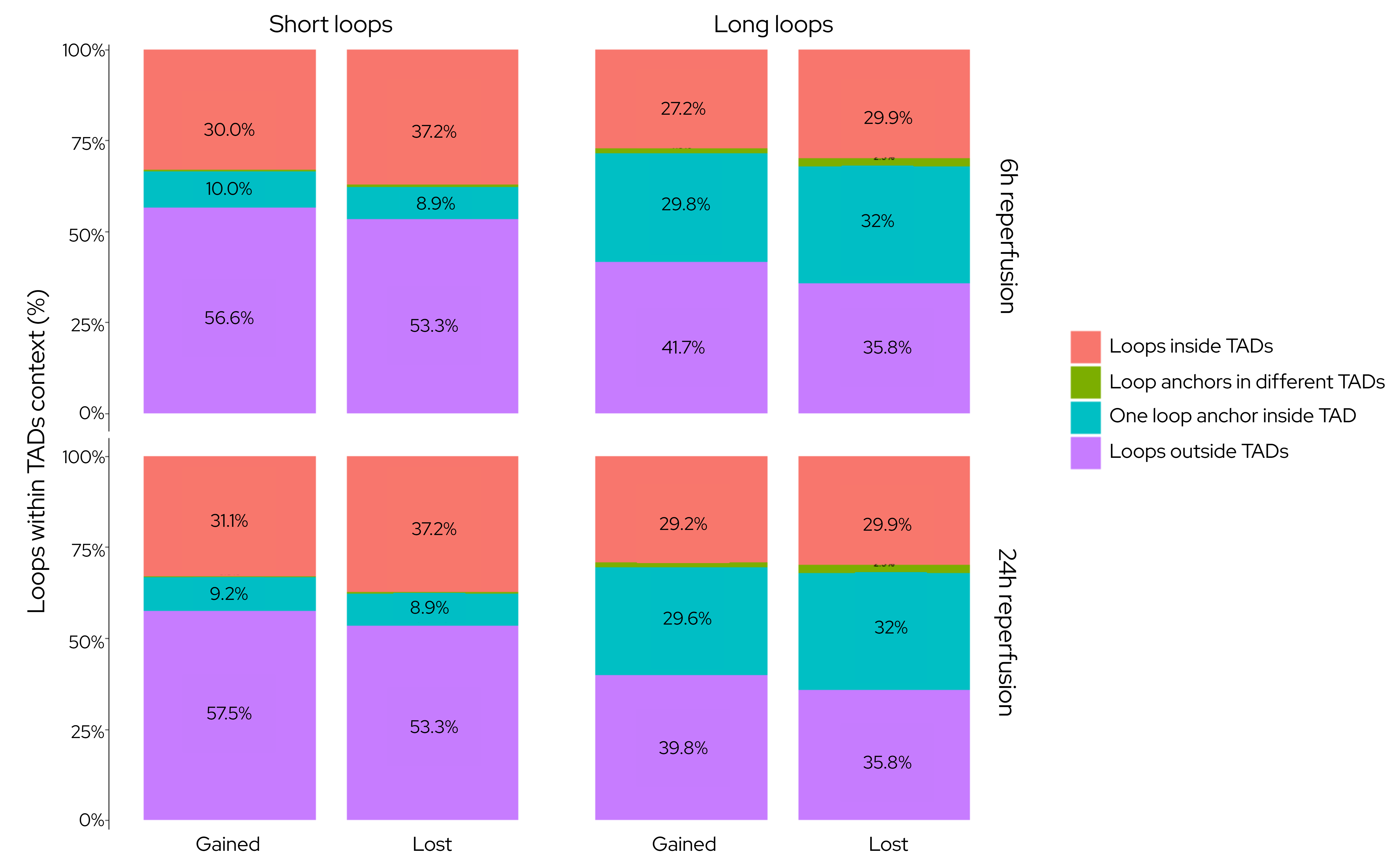

**Figure S10:** Spatial distribution of dynamic chromatin loops relative to TAD contexts. Chromatin loops were classified by length (Short vs. Long) and by their dynamics (Gained vs. Lost) following 6 h and 24 h of reperfusion. Loops were further categorized based on their relationship to Topologically Associating Domains (TADs): loops fully contained within a TAD (red), loops with anchors spanning different TADs (green), loops with one anchor inside a TAD (teal), and loops entirely outside TAD boundaries (purple). Values are expressed as the percentage of total loops within each category, enabling direct comparison between gained and lost features.

M6h - short

M24h - short
